# Multiomics-based assessment of the impact of airflow on diverse plant callus cultures

**DOI:** 10.1101/2024.07.17.604000

**Authors:** June-Sik Kim, Muneo Sato, Mikiko Kojima, Muchamad Imam Asrori, Yukiko Uehara-Yamaguchi, Yumiko Takebayashi, Thi Nhung Do, Thi Yen Do, Kieu Oanh Nguyen Thi, Hitoshi Sakakibara, Keiichi Mochida, Shijiro Ogita, Masami Yokota Hirai

## Abstract

Plant cell culture has multiple applications in biotechnology and horticulture, from plant propagation to the production of high-value biomolecules. However, the interplay between cellular diversity and ambient conditions influences the metabolism of cultured tissues; understanding these factors in detail will inform efforts to optimize culture conditions. This study presents multiomics datasets from callus cultures of tobacco (*Nicotiana tabacum*), rice (*Oryza sativa*), and two bamboo species (*Phyllostachys nigra* and *P. bambusoides*). Over four weeks, calli were cultured under continuous moisture without airflow or gradually reduced ambient moisture with airflow. For each sample, gene expression was profiled with high-throughput RNA sequencing, 442 metabolites were measured using liquid chromatography (LC) with triple-quadrupole mass spectrometry (LC–QqQMS), and 31 phytohormones were quantified using ultra-performance LC coupled with a tandem quadrupole mass spectrometer equipped with an electrospray interface (UPLC-ESI-qMS/MS) and ultra-high-performance LC– orbitrap MS (UHPLC-Orbitrap MS). These datasets highlight the impact of airflow on callus cultures, revealing differences between and within species, and provide a comprehensive resource to explore the physiology of callus growth.

## Background & Summary

Plant cell cultures, historically used for conservation and molecular breeding, are now seen as potential cell factories for sustainable plant-based production of high-value biomaterials including pharmaceuticals and cosmetics, facilitated by genetic engineering and advanced by synthetic biology ^1–3^. Using plant cell cultures as cell factories for production of biomaterials requires a deep understanding of the intricate biological systems in cultured cells. Indeed, plant callus cultures derived from various plant species exhibit remarkable diversity in their metabolic signatures ^4^, as well as in their capacities for proliferation, regeneration, and differentiation ^5^. Such diversity in callus cultures is often observed between plant species and even within the same species, influenced by various factors including the origin of the tissues ^6^, plant genotype ^7^, and the ambient conditions ^8^. Characterizing the cellular states and responses is crucial not only for illustrating the inherent metabolic potential of plant species but also for optimizing culture conditions, guiding genetic modifications, and predicting metabolic outcomes.

In characterizing the cellular states and responses of callus cultures, a multiomics approach provides invaluable datasets that capture multifaceted snapshots of cellular systems. The integration of transcriptomic and metabolomic data enables us not only to assess cellular states and responses but also to explore potential key regulatory factors that may govern metabolic pathways. Phytohormones orchestrate virtually every aspect of plant growth, development, differentiation, and responses to environmental conditions. As the proliferation and differentiation abilities of plant cells are also regulated by the interactions of phytohormones, the simultaneous profiling of these key signals provides important information on the cellular state. The collective impact of such integrated and comparative omics insights facilitates the selection and modification of the cellular chassis, harnessing the full potential of metabolic diversity in plant cells for the development of efficient cell factories.

Here, we generated multiomics datasets encompassing transcriptome, hormonome, and metabolome data from cultured calli of tobacco (*Nicotiana tabacum*), rice (*Oryza sativa*), and two bamboo species (*Phyllostachys nigra* and *P. bambusoides*) (Fig. 1a), measured over four weeks. The calli were grown under conditions that either maintained continuous moisture without airflow or gradually reduced the ambient moisture with airflow, determined by the use of different types of Petri dish seals (Table S1; Fig. 1b and c). The dataset includes 24 samples per species (three replicates × four timepoints × two airflow conditions), which were analyzed to explore how the different airflow conditions affect their cellular states. For our transcriptome analysis, we constructed Illumina-compatible RNA-sequencing (RNA-seq) libraries and conducted 150-bp paired-end sequencing (Fig. 1d), obtaining more than 25.2 million high-quality reads per sample. Our metabolome profiling quantified 442 distinct metabolites using a widely targeted metabolomics platform with liquid chromatography and triple-quadrupole mass spectrometry (LC-QqQMS). Moreover, 31 phytohormones were quantified using both ultra-performance LC (UPLC) coupled with a tandem quadrupole mass spectrometer (qMS/MS) equipped with an electrospray interface (ESI) (UPLC-ESI-qMS/MS) and ultra-high-performance LC–orbitrap high-resolution MS (UHPLC-Orbitrap MS), providing comprehensive profiles of these phytohormones. This multiomics dataset, the first to systematically capture data across diverse plant species’ callus cultures, provides foundational insights into the cellular states and responses of calli, enhancing our understanding of plant cellular physiology and supporting future efforts in metabolic engineering for efficient plant-based material production.

**Figure 1.**
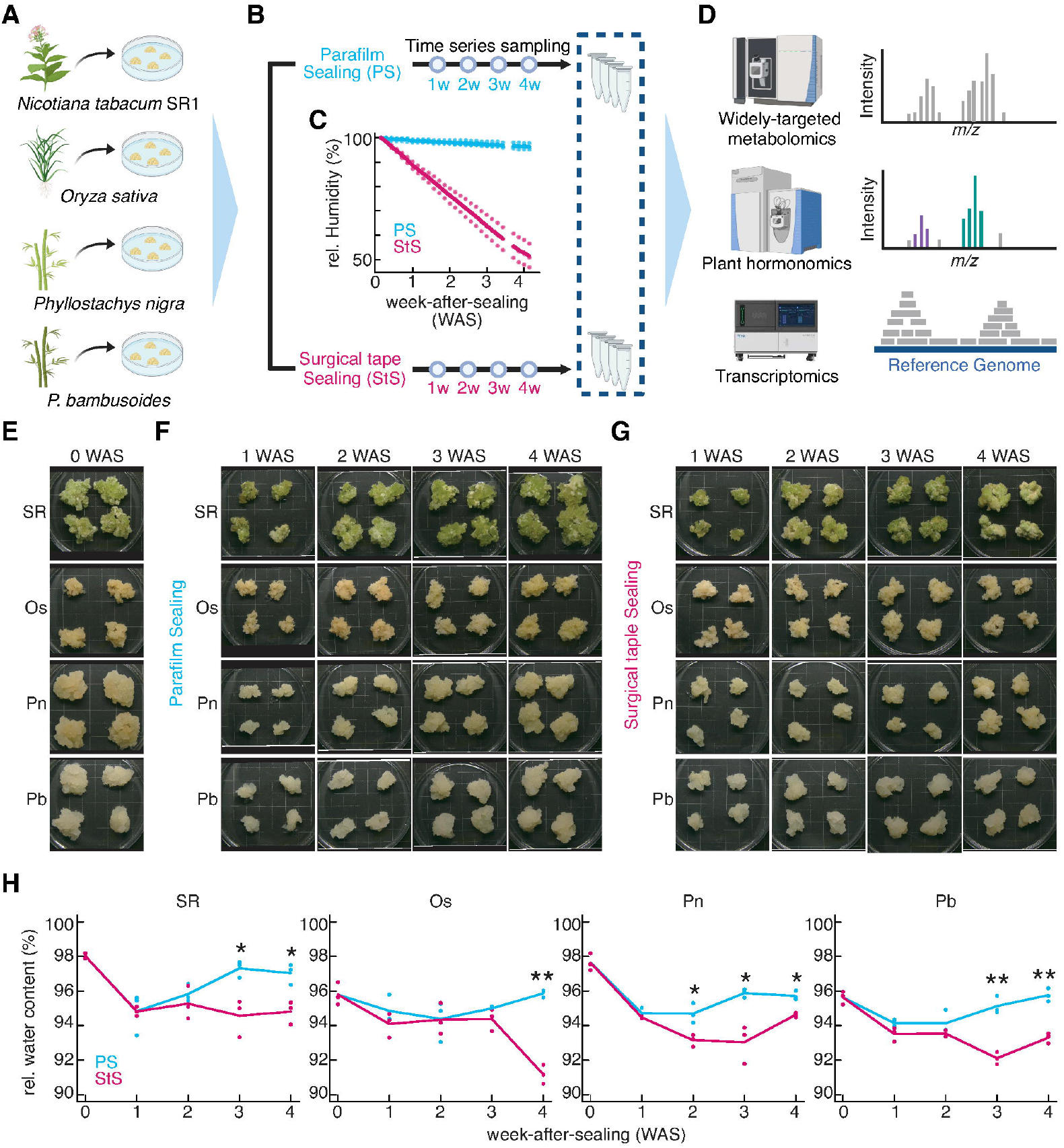
Experimental scheme and callus samples. **(A)** Callus induction from tobacco (*Nicotiana tabacum* cv. SR1 [SR]), rice (*Oryza sativa* cv. Nipponbare [Os]), and two bamboo species (*Phyllostachys nigra* [Pn] and *P. bambusoides* [Pb]). **(B)** Humidity perturbation: Parafilm sealing (PS) for humid conditions and surgical tape sealing (StS) for dehydration conditions. Cultures were maintained for four weeks and sampled weekly. **(C)** Humidity was monitored by weight changes in the Petri dishes after sealing (n = 3). Data points were collected daily. **(D)** Multiomics analysis of the callus samples. For the transcriptome analysis, Illumina-compatible libraries were prepared, and stranded paired-end RNA-seq reads were obtained. The RNA-seq reads were mapped to the respective reference genomes, and gene expression levels were estimated based on read counts. The metabolome analyses was conducted using LC-QqQMS, and the hormone analysis was conducted using UPLC-ESI-qMS/MS and UHPLC-Orbitrap MS. The metabolome analysis quantified 442 compounds, while the hormone analysis quantified 31 hormones. **(E–G)** Visual appearance of callus samples. **(E)** Initial appearance under PS conditions before subculture. **(F–G)** Time-series visual appearance under PS conditions **(F)** and StS conditions **(G)** over four weeks of subculturing. The background grids are 10 mm × 10 mm in size. **(H)** Relative water content changes in callus samples after four weeks of subculturing under PS and StS conditions. Statistically significant differences are indicated as \*\**p* < 0.05 and \**p* < 0.01 (two-sided Student’s t-test). Some figures in **(A–D)** were generated using BioRender.

## Methods

### Growth conditions

The tobacco callus used in this study was induced from leaf segments of an SR1 plant (*Nicotiana tabacum* L. cv. SR1 [SR]), using seeds provided by the Leaf Tobacco Research Center (Japan Tobacco, Tokyo, Japan). The SR callus was cultured on Linsmaier and Skoog medium (pH 5.7) containing 1 μM 2,4-D, 3% sucrose, and 0.3% gellan gum in sterilized Petri dishes with a diameter of 90 mm and a height of 15 mm. The calli of rice (*Oryza sativa* L. cv. Nipponbare [Os]) and two bamboo species (*Phyllostachys nigra* (Lodd. ex Lindl.) Munro var. Henonis [Pn] and *P. bambusoides* Siebold and Zucc. [Pb]) were cultured on a modified Murashige and Skoog medium (pH 5.7) containing 10 μM picloram, 3% sucrose, and 0.3% gellan gum in sterilized Petri dishes of the same dimensions. The calli derived from these four species were routinely subcultured by transferring four pieces of callus, each approximately 500 mg in fresh weight and 10 mm in diameter, to a new Petri dish with fresh medium every four weeks. The SR and Os callus cultures were maintained at 25°C with a 16-h light:8-h dark photoperiod, with light provided by fluorescent illumination (65 μmol m^−2^ s^−1^), while the bamboo callus cultures were maintained at 25°C in the dark ^9^.

The different humidity conditions were achieved by sealing the Petri dishes either with Parafilm M (Amcor, Victoria, Australia) for the non-airflow condition (Parafilm sealing; PS) or with Micropore Surgical Tape (3M, Maplewood, Minnesota, USA) for the airflow condition (surgical tape sealing; StS) (Fig. 1b). The weights of the Petri dishes were measured every day for four weeks before culturing the calli to determine the changes in their humidity levels caused by the different sealing methods (Fig. 1e-1g). For the comparison of the water content within the callus, the relative water content was determined by measuring the fresh weight and dry weight of the sampled callus and calculating the difference (Fig. 1h). After subculturing, callus samples were taken weekly for four weeks (Table S1).

Each callus sample was divided into two pieces. One was frozen in liquid nitrogen for the transcriptome analysis, while the other was freeze-dried for the metabolome and hormone analyses. The samples were stored at –80°C until the analyses were performed.

### Metabolome analysis

A 4-mg dry-weight portion of each sample was used for the analysis. Metabolite extraction was performed according to the method described previously ^10^. A widely targeted metabolome analysis was conducted as described previously ^11^, using the selective reaction monitoring (SRM) conditions of 442 standard metabolites (Table S2). The peak area of each metabolite was calculated using MRMPROBS version 2.60 ^12,13^.

### Hormone analysis

A 20-mg dry-weight portion of each sample was used for the analysis. The extraction and quantification of plant hormones were performed as described previously ^14,15^. Endogenous cytokinins were measured using UPLC and Octadecylsilyl (ODS) columns (AQUITY Premier HSS T3, 1.8 μm, 2.1 × 100 mm; Waters, Milford, Massachusetts, USA) combined with a qMS/MS equipped with an ESI (UPLC-Xevo TQ-XS; Waters). The data were processed using MASSLYNX with TARGETLYNX version 4.2 (Waters). Endogenous gibberellins, salicylic acid, abscisic acid, jasmonic acid, jasmonoyl isoleucine, and auxins were measured using UHPLC and ODS columns (AQUITY Premier HSS T3, 1.8 μm, 2.1 × 100 mm; Waters) combined with Q-Exactive (Thermo Fisher Scientific, Waltham, Massachusetts, USA) without MS probe modification. The data were processed using XCALIBUR version 4.5 (Thermo Fisher Scientific).

### Transcriptome analysis

The same samples used for the metabolome and hormonome analyses were also subjected to a next-generation sequencing–aided transcriptome analysis of biological triplicate samples. Total RNA was extracted from each homogenized sample using an ISOSPIN Plant RNA kit (Nippon Gene, Tokyo, Japan). The mRNA was isolated and the sequencing library was prepared using NEXTFLEX Poly(A) Beads 2.0 (PerkinElmer, Waltham, Massachusetts, USA) and a NEXTFLEX Rapid Directional RNA-Seq Kit 2.0 (PerkinElmer), respectively. Quality checks and the quantification of the prepared libraries were performed using a TapeStation system (Agilent Technologies, Santa Clara, California, USA).

Sequencing on a DNBSEQ-G400 (MGI Tech, Shenzhen, China) system yielded an average of 25.2 million paired-end reads (2 × 150 bp) per library. The read quality was assessed using FastQC version 0.12.1 ^16^, and the subsequent steps for filtering artificial and low-quality sequences were performed using Trimmomatic version 0.39 ^17^ with the following parameters: “ILLUMINACLIP:2:30:10 LEADING:20 TRAILING:20 SLIDINGWINDOW:4:15 MINLEN:36”. Three reference genomes were adopted as matrices for mapping the reads from different sample origins: Nitab4.5 (https://www.ncbi.nlm.nih.gov/datasets/genome/GCA_002210045.1/) ^18^ for SR, IRGSP-1.0 (https://rapdb.dna.affrc.go.jp/download/irgsp1.html) ^19^ for Os, and a chromosome-level genome assembly of diploid moso bamboo (*P. edulis*) (http://gigadb.org/dataset/view/id/100498) ^20^ for Pn and Pb. The expression levels of these bamboo species were estimated based on loci annotated in the *P. edulis* genome through cross-species mapping, which does not account for the homoeologous gene duplications or unique genomic features of Pn and Pb.

The cleaned reads were aligned to the allocated reference matrix using the two-pass mode of STAR version 2.7.10a ^21^ with default parameters for rice and bamboo reads, while applying the following parameters for tobacco reads: “--outFilterScoreMinOverLread 0 -- outFilterMatchNminOverLread 0.15 --outFilterMatchNmin 0.15”, for relaxed mapping capacity. The digital expression value for each genetic locus was estimated using RSEM version 1.3.1 ^22^ by applying the STAR-generated.bam file as an input. The overview of the transcriptome data and their visualization were achieved using R version 4.3.2 ^23^. The tximport version 1.28 ^24^ was employed to incorporate and combine the STAR-RSEM resultants into gene expression matrices.

## Data Records

### Metabolome and hormonome data

The LC-MS/MS raw datasets from our widely targeted metabolome analysis are available from Data Resources Of Plant Metabolomics (DROP Met) under the accession number DM0060 on the Platform for RIKEN Metabolomics (PRIMe) website (https://prime.psc.riken.jp/menta.cgi/prime/drop_index).

### Transcriptome data

Raw RNA-seq read data derived from the callus samples of tobacco (SR), rice (Os), and the two bamboo species (Pn and Pb) are available from the DNA Database of Japan (DDBJ) BioProject no. PRJDB18353. The processed data have been deposited in the DDBJ Genomic Expression Archive with the specific accession numbers E-GEAD-653 (SR), E-GEAD-652 (Os), E-GEAD-654 (Pn), and E-GEAD-655 (Pb).

## Technical validation

### Metabolome analysis

For the widely targeted metabolome analysis, internal standard compounds (8.4 nm lidocaine for positive-ion mode and 210 nm 10-camphorsulfonic acid for negative-ion mode) were added to the extraction solvent to normalize the metabolite signals. The quality-control sample, a mixture of equal amounts of all samples, was analyzed once in every ten samples to check for significant fluctuations in signal intensity. After automated peak picking by MRMPROBS version 2.60 ^12,13^, visual confirmation was performed to adjust the baseline and verify the peak shape before calculating the peak area.

### Hormonome analysis

To measure the phytohormones, stable isotope-labeled internal standards of almost all target phytohormones were added to the extraction solvent to allow the accurate correction of losses resulting from the pre-treatment of the samples, such as solid-phase extraction. In the LC–MS measurements, target molecular species could be more accurately measured by separation using the multiple reaction monitoring (MRM) method (Xevo TQ-XS; Waters) or the selected ion monitoring (SIM) method with high-resolution accurate mass (Q-Exactive; Thermo Fisher Scientific). As the retention time of a target molecular species may vary depending on the circumstances, the retention times were confirmed by measuring a reference material once every 24 samples during the measurement. After automated peak picking by the analytical software, visual confirmation was performed to adjust the baseline and verify the peak shape. The standard curve had a squared Pearson’s correlation coefficient (r^2^) value of at least 0.99 and any peaks outside the range of the standard curve were excluded.

### Transcriptome analysis

#### Quality control of the raw RNA-seq reads and filtering

The read statistics and results of the quality control for the RNA-seq data are detailed in Table S3. Based on the FastQC results, the proportion of reads with a mean Phred quality score of 30 (Q30) or higher was an average of 95.58% ± 1.32%. After removing artificial and low-quality (less than Q30) sequences, the proportion of clean and properly paired reads to raw reads was an average of 96.95% ± 3.97%, with an average read length of 148.67-nt, indicating a high percentage of qualified reads in our datasets.

#### Clean reads mapping to the reference sequences

The statistics from mapping the clean reads of each sample to their allocated reference genomes are summarized in Table S4. The average proportions of properly paired mapped reads to the input reads were 88.73% ± 0.24% for SR, 94.78% ± 0.52% for Os, and 90.17% ± 0.39% and 90.93% ± 0.27% for the two bamboo species, respectively. These mapped reads covered 20,956 ± 464 loci of the whole 35,519 protein-coding tobacco genes in SR, 27,850 ± 651 loci of the 37,850 rice genes in Os, and 29,797 ± 471 and 32,859 ± 684 loci of the 50,397 genes annotated in the *P. edulis* genome detected in Pn and Pb, respectively. Furthermore, by counting the loci with significant expression levels (≥1 transcripts per million, TPM), the evaluated SR data covered 52.3% ± 0.97% of the tobacco genes, the Os data covered 56.9% ± 1.48% of the rice genes, and the Pb and Pn data respectively covered 48.8% ± 0.32% and 53.2% ± 1.07% of the bamboo genes. While coverage ratios varied by different integrities of the allocated reference genomes, these results indicate that the RNA-seq samples possess both the quality and quantity of reads required to obtain reliable gene expression profiles.

### Reproducibility of the biological replicates

To assess the reproducibility among biological replicates, we assessed Pearson’s correlation coefficients (PCCs) across the samples. The average within-replication PCCs of the transcriptome TPM data of significantly expressed genes (≥1 TPM) were 0.93 ± 0.03 for SR, 0.95 ± 0.03 for Os, 0.96 ± 0.05 for Pn, and 0.97 ± 0.01 for Pb within replications (Fig. 2). The within-replication PCCs of the merged abundance set of metabolome and hormonome data were as follows: 1.00 ± 0.00 for SR, 0.97 ± 0.03 for Os, 0.99 ± 0.01 for Pn, and 0.99 ± 0.01 for Pb (Fig. 2). These high levels of correlation strongly support the reliability of our dataset, based on the high reproducibility of every procedure of sampling, data acquisition, and data processing.

**Figure 2.**
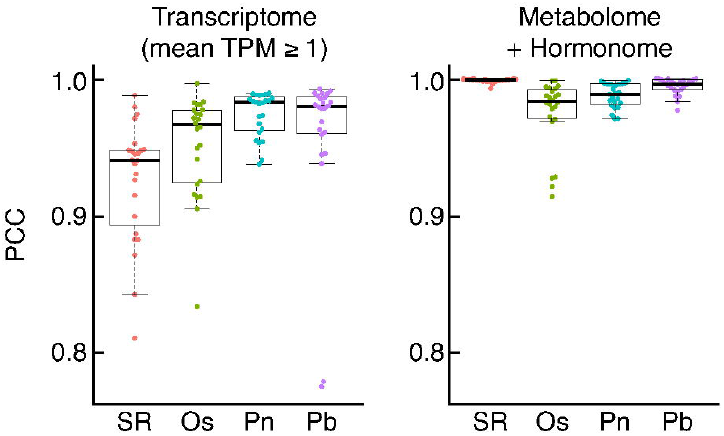
Reproducibility of biological replicates. The distribution of Pearson correlation coefficients (PCCs) for the biological replicates. The left panel displays the profiles of the transcriptomes (genes with mean TPM ≥ 1) of each species while the right panel displays those of the metabolome and hormonome. SR: *Nicotiana tabacum* cv. SR1, Os: *Oryza sativa* cv. Nipponbare, Pn: *Phyllostachys nigra*, and Pb: *P. bambusoides*.

## Usage Notes

The widely targeted metabolome and hormonome analyses enable the quantification of metabolites and hormones across plant species, allowing the diversity in cellular states among species to be represented in a unified space. Fig. 3 shows the principal component analysis (PCA) plots of callus samples from the four plant species used in this study, representing the comparative distribution of samples based on metabolite and hormone accumulation. As expected, the sample distribution shows distinct species-specific clustering, with the two bamboo species (Pn and Pb) positioned closely together. As shown in Fig. 1H, the relative water content of these calli significantly decreased after four weeks of culture in Petri dishes sealed with surgical tape for all these plant species. The patterns of response to this decrease in water content, as observed in the metabolome and hormonome data, varied among species; for example, tobacco showed significant changes in the PCA space, while the other samples from the grass family were plotted close to each of their control conditions (Fig. 3A).

**Figure 3.**
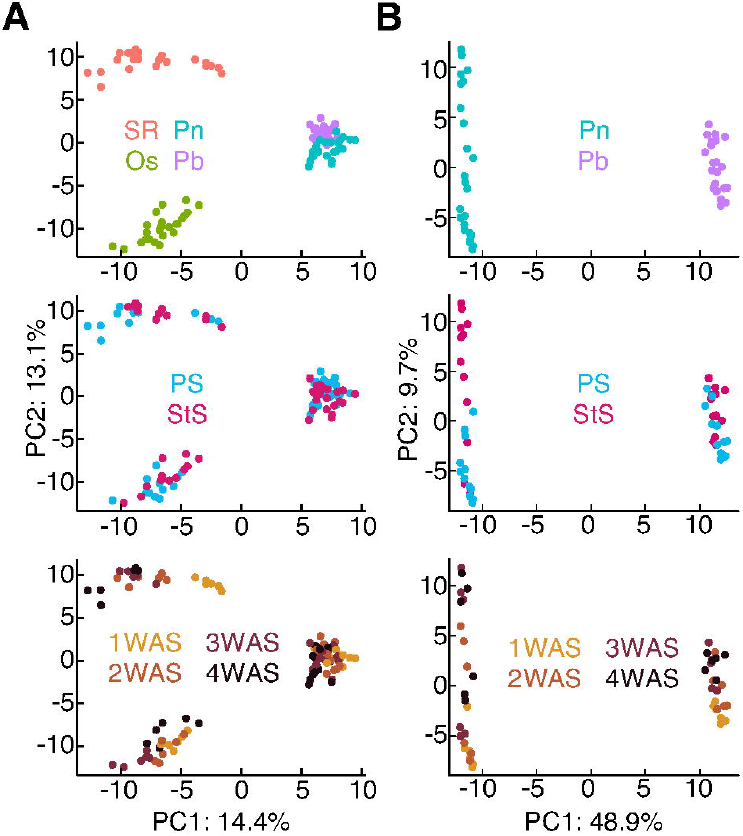
Metabolomic and hormone profiles. **(A)** Principal component analysis (PCA) plots of the combined metabolome and hormone data from the callus cultures derived from the four plant species. SR: *Nicotiana tabacum* cv. SR1, Os: *Oryza sativa* cv. Nipponbare, Pn: *Phyllostachys nigra*, and Pb: *P. bambusoides*, represented with properties of species (upper panel), humidity conditions (middle panel), and sampling time (lower panel). **(B)** PCA plots of the combined metabolome and hormone data from the callus cultures derived from Pn and Pb, represented with the properties of species, humidity conditions, and sampling time.

Although the callus samples of the two bamboo species were closely clustered in Fig. 3, a separate PCA focusing only on these two species distinctly visualizes their physiological states and responses to different humidity conditions. Comparing the responses of callus samples to humidity differences between the two bamboo species indicates that *P. nigra* exhibits a greater magnitude of response to the decreased water content than *P. bambusoides* (Fig. 3B). These results demonstrate that the physiological states of plant callus cells and their responses to decreased humidity and subsequent reduction in water content are diverse, not only among distantly related species but also within the same genus. We highlight the importance of carefully assessing the adaptability of callus cells in response to various culture conditions to establish chassis strains for robust and scalable plant cell factories. The datasets provided in this study represent the first systematically collected multiomics dataset for various plant callus samples, offering foundational comparative insights that facilitate the development of cell factories harnessing the metabolic diversity of plants.

## Code Availability

Detailed commands for the transcriptome analysis and in-house R codes for data visualization used in this study are available through a GitHub repository (https://github.com/junesk9).

## Acknowledgments

This research was supported by the RIKEN Cluster for Science, Technology and Innovation Hub (RCSTI) and a GteX Program Japan Grant (no. JPMJGX23B0) to K.M. and M.Y.-H.

## Supplementary information

**Table S1**. Sample details

**Table S2**. Compounds measured in the widely targeted metabolome analysis and their conditions for multiple reaction monitoring (MRM) profiling in this study

**Table S3**. RNA-seq reads statistics and quality control

**Table S4**. RNA-seq mapping result statistics

